# High-Concordance Validation of Droplet Digital PCR and Next-Generation Sequencing for EGFR Mutation Detection Across Diverse Biospecimens in a Large-scale NSCLC Cohort Study

**DOI:** 10.1101/2025.11.05.686693

**Authors:** Ming Liu, Linping Lu, Lingxiang Zhu, Xiaoni Zhang, Yuanyuan Liu, Xiaojing Ren, Shimin Liu, Shaochen Cheng, Mingyan Xu, Chao Lu, Yan Peng, Wenmei Su, Yong Guo, Shifu Chen

## Abstract

**Backgrounds:** Next-generation sequencing (NGS) and droplet digital PCR (ddPCR) are both established methods for detecting EGFR mutations in non-small cell lung cancer (NSCLC). However, comprehensive validation of their concordance in mutation detection and variant allele frequency (VAF) quantification across heterogeneous sample types remains limited. Inconsistent results from different sample types (e.g., cfDNA, FFPE) pose a significant challenge to clinical decisions. This is particularly critical for advanced NSCLC patients, who often rely on liquid biopsy, yet the concordance between liquid and tissue-based testing lacks validation in large-scale studies. Another issue in clinical practice is that many tumor samples are limited in quantity, making it difficult to meet testing requirements. Therefore, it is worth exploring whether pre-capture NGS libraries can serve as substitutes for original DNA.

**Methods:** In this study, we first developed three assays to detect EGFR L858R, exon 19 deletions (Ex19del) and T790M, respectively using ddPCR platform. Their Limit of Detection (LOD) could reach 0.01% at 100 ng of input DNA. Subsequently, we conducted a large retrospective clinical study to systematically compare the detection performance of ddPCR and NGS across three mutation types using approximately 1,000 EGFR-positive samples, including cell-free DNA (cfDNA), pre-capture NGS libraries of cfDNA (cfDNA-prePCR), FFPE-derived DNA (ffpeDNA), pre-capture NGS libraries of FFPE-derived DNA (ffpeDNA-prePCR), fresh tumor tissue DNA (ttDNA), pre-capture NGS libraries of ttDNA (ttDNA-prePCR), pleural effusion supernatants DNA (peDNA), and pre-capture NGS libraries of peDNA (peDNA-prePCR). They were analyzed for detection concordance and VAF correlation. Especially, we made comparisons of the tumor DNA, including ctDNA and tumor tissue DNA, with their paired pre-capture NGS library.

**Results:** Key findings demonstrated excellent overall agreement between NGS and ddPCR. The mutation detection concordance rates were 98.72% (overall), with subtype-specific rates of 98.93% (L858R), 99.23% (Ex19del), and 97.14% (T790M). VAF measurements between ddPCR and NGS showed exceptional correlation (Pearson’s r = 0.975, P<0.001). Notably, pre-capture NGS libraries showed remarkable VAF concordance with their source materials (0.993 for cfDNA libraries vs cfDNA; 0.998 for tumor tissue libraries vs tumor DNA with EGFR L858R; 0.991 for tumor tissue libraries vs tumor DNA with EGFR Ex19del).

**Conclusions:** NGS and ddPCR demonstrate high concordance in EGFR mutation detection and VAF quantification, supporting their complementary roles in clinical testing. Using pre-capture libraries as an alternative to source samples can avoid repeat biopsies and enables subsequent testing for patients with inadequate FFPE sample quantity. These findings establish an evidence base for integrated diagnostic paradigms leveraging NGS’s multiplexing power and ddPCR’s sensitivity.

## Introduction

In the management of lung cancer, EGFR L858R and exon 19 deletions (Ex19del) serve as primary targets for first-line tyrosine kinase inhibitors (TKIs) **[1]**, while the T790M mutation determines eligibility for third-generation TKI therapy. **[2]** Collectively, these three alterations account for over 90% of clinically relevant EGFR mutations **[3]**, making their accurate detection critical for treatment decision-making.

Current clinical practice presents several diagnostic challenges. While tissue samples remain the gold standard for initial diagnosis **[4]**, plasma or pleural effusion samples often become necessary for advanced or treatment-resistant cases. **[5]** However, the reliability of mutation detection across these diverse sample types lacks comprehensive validation. The two predominant detection methods - next-generation sequencing (NGS) and droplet digital PCR (ddPCR) - each present unique technical considerations. **[6, 7]** For Ex19del, a deletion mutation, the potential amplification biases during NGS library preparation may introduce variability in variant allele frequency (VAF) quantification. **[8]** Although L858R and T790M (point mutations) demonstrate greater detection stability, their accuracy in low-VAF samples remains debated. **[9]** Furthermore, the clinical application scenarios for NGS (with its multi-gene profiling advantage) versus ddPCR (with superior single-target precision) require clearer definition. **[10]**

Existing studies face significant limitations, including small sample sizes and restricted sample type representation **[11, 12]**, thereby offering limited clinical guidance. To address these gaps, our study systematically evaluated the concordance between NGS and ddPCR in VAF measurement for EGFR L858R/Ex19del/T790M mutations across nearly 1000 lung cancer cases, encompassing plasma, FFPE tissues, fresh tumor tissues, pleural effusion supernatants, and plasma cfDNA NGS libraries. Notably, we performed focused validation of VAF concordance between ddPCR and NGS libraries for both plasma cfDNA and tumor tissue DNA, specifically examining L858R (point mutation) and Ex19del (deletion mutation). This comprehensive analysis provides crucial evidence for assessing NGS reliability and optimizing clinical testing strategies for these therapeutically pivotal EGFR mutations.

## Materials and Methods

### Study Population and Sample Characteristics

This retrospective clinical investigation enrolled 789 patients with pathologically confirmed non-small cell lung cancer (NSCLC). The study data were collected between January 1, 2023 and February 5, 2025, with all clinical specimens and corresponding data obtained from Shenzhen HaploX Medical Laboratory (Shenzhen, China). The study protocol was conducted in strict accordance with medical ethics guidelines and was reviewed and approved by the Institutional Review Board of The Third Affiliated Hospital of Shenzhen University. Written informed consent was obtained from all participating patients prior to treatment initiation. The inclusion criteria comprised: (1) pathologically confirmed NSCLC diagnosis; (2) identification of EGFR mutations (L858R, Ex19del, or T790M) through initial next-generation sequencing (NGS) screening; and (3) availability of complete clinical documentation, including disease staging, treatment history, sample type specifications, and collection timelines. The final cohort included 952 cases.

### Sample Processing and Nucleic Acid Extraction

Peripheral blood samples were collected using EDTA-containing tubes and processed within two hours of collection. Initial centrifugation was performed at 1,600 × g for 10 minutes at 4°C to separate plasma from cellular components. The supernatants were further centrifuged at 10,000 g for 10 min at 4 °C, and plasma was harvested., while the peripheral blood lymphocyte (PBL) pellet was retained for genomic DNA extraction.

Circulating cell-free DNA (cfDNA) was isolated from a minimum of 2 mL plasma using the QIAamp Circulating Nucleic Acid Kit (QIAGEN) according to the manufacturer’s specifications. Parallel genomic DNA (gDNA) extraction from PBLs was conducted using the RelaxGene Blood DNA System (TianGen Biotech Co., Ltd., China) to serve as matched normal controls. DNA quantification was performed using the Qubit 2.0 Fluorometer (Thermo Fisher Scientific, USA) with appropriate dsDNA HS Assay Kits, following manufacturer-recommended protocols. The integrity and size distribution of extracted plasma cfDNA were evaluated through fragment analysis using the Agilent 2100 Bioanalyzer System with the High Sensitivity DNA Kit. The quality control criteria included the presence of a characteristic nucleosomal fragmentation pattern with a dominant peak at approximately 160 bp, consistent with mononucleosomal DNA fragments.

FFPE tissue sections underwent xylene-based deparaffinization through three sequential washes, followed by ethanol rehydration. DNA extraction was performed using the QIAamp DNA FFPE Tissue Kit (QIAGEN) with optimized protocols for degraded samples. DNA integrity was assessed through degradation analysis, with all samples required to meet a minimum DV200 value of 30% (percentage of DNA fragments >200 bp).

Fresh tissue specimens were mechanically homogenized using the TissueLyser II system (QIAGEN) with appropriate lysing matrix tubes. Subsequent DNA extraction was carried out with the QIAamp DNA Mini Kit (QIAGEN) following standard protocols. DNA concentration and purity were determined via Qubit quantification.

Pleural effusion samples underwent initial cellular component removal through centrifugation at 3,000 × g for 15 minutes. The resulting supernatant was subjected to additional filtration through 0.8 μm membranes to minimize potential matrix interference. Pleural effusion-derived DNA (peDNA) extraction followed the identical protocol established for plasma cfDNA isolation. Final DNA quantification was performed using the Qubit 2.0 system according to manufacturer specifications.

All extracted DNA samples were aliquoted and stored at -80°C until subsequent analysis to maintain nucleic acid integrity and prevent degradation.

### Detection Methods

#### Next-Generation Sequencing (NGS) Methodology

##### Library Construction

DNA extracted from tumor tissues, including FFPE and fresh tumor tissues and matched germline samples underwent enzymatic fragmentation using dsDNA Fragmentase (New England Biolabs). DNA fragments ranging from 150-250 base pairs were size-selected using AMPure XP beads (Beckman Coulter, Inc., Brea, CA, USA). Library preparation was performed using the KAPA Library Preparation Kit (Kapa Biosystems, Inc., Wilmington, MA, USA) according to manufacturer’s specifications. cfDNA and peDNA underwent direct library preparation. The protocol included initial quality assessment, end repair, 3’-end A-tailing, and adapter ligation conducted at 20°C for 15 minutes. Dual-SPRI size selection was subsequently performed to enrich appropriately ligated fragments. Amplification of ligated fragments was carried out in 50 μL reactions using 1× KAPA HiFi Hot Start Ready Mix with Pre-LM-PCR Oligos, with PCR cycles (7-12 cycles) optimized according to input DNA quantity. The thermal cycling conditions consisted of: initial denaturation at 98°C for 45 seconds (1 cycle); denaturation at 98°C for 15 seconds, annealing at 60°C for 30 seconds, and extension at 72°C for 30 seconds (7-12 cycles); final extension at 72°C for 1 minute (1 cycle). Library concentration was quantified using the Qubit dsDNA HS Assay kit, and fragment size distribution was verified with the DNA 1000 Kit on the Agilent 4200 Bioanalyzer system.

##### Targeted Capture and Sequencing

Targeted capture was performed using a custom-designed 1326 gene panel (HaploX Biotechnology, Shenzhen, China), an sequencing panel with comprehensive coverage of EGFR exons 18-21. Hybridization capture was conducted according to manufacturer’s protocols for 16-20 hours at 47°C. Post-capture amplification was performed using 12-14 PCR cycles with 1× KAPA HiFi Hot Start Ready Mix under identical thermal cycling conditions as library construction. Reactions were pooled and purified using AMPure XP beads. Final libraries were sequenced with 150 bp paired-end reads on the Illumina NovaSeq 6000 platform (Illumina, San Diego, CA, USA), achieving minimum sequencing depths of 1,000× for tissue specimens and 2,000× for liquid biopsy samples.

#### Bioinformatic Analysis Pipeline

##### Raw Data Processing and Alignment

Raw sequencing data underwent quality control and adapter trimming using fastp (v0.12.6). Processing included removal of reads containing N bases exceeding 5 bp threshold, elimination of reads with low-quality bases (quality score ≤20) comprising >40% of read length, and sliding window trimming. Quality-filtered reads were aligned to the human reference genome (hg19/GRCh37) using Burrows-Wheeler Aligner (BWA v0.7.15) with default parameters. PCR duplicates were removed using Gencore (v0.12.0), and mpileup files were generated using Samtools (v0.1.19) for properly paired reads with mapping quality ≥60.

##### Variant Calling, Filtering, and Annotation

Single nucleotide variants (SNVs) and short insertions/deletions (indels) were identified by VarScan2 v2.3.8 [13]. The average sequencing depth after deduplication in tumor tissues DNA is ≥500×, and the average sequencing depth after deduplication of cfDNA is ≥1500×. For cfDNA, the filter criteria were set as follows: The somatic variants (SNVs or indels) presented at least 5 unique reads, at least 1 on each strand, and less than 0.5% mutant allelic frequency in the paired normal sample (PBLs) were retained. Additionally, for cfDNA, we excluded any SNVs and indels by the background polishing using cfDNA samples from healthy subjects. For tumor tissue DNA, the filter criteria were set as follows: The somatic variants (SNVs or indels) presented at least 5 unique reads, at least 1 on each strand, and less than 0.5% mutant allelic frequency in the paired normal sample (PBLs) were retained. A manual visual inspection step was applied to further remove artifacts by GenomeBrowse [14]. All SNVs/indels were annotated using ANNOVAR v2018-04-16 [15].

### Droplet Digital PCR (ddPCR) Methodology

#### Assay Design and Experimental Configuration

Independent ddPCR reactions were designed to detect the presence of EGFR L858R (c.2573T>G, 2573_2574TG>GT), EGFR Ex19del and EGFR T790M (c.2369C>T).

The L858R and T790M mutations were detected using specific probes, while the Ex19del was detected using specific primers. The primers and probes for dPCR were shown in **Table 1**. The primers 19del-F/19del-R and the HEX-labeled EGFR-WT probe were specifically designed to ensure that none of the oligonucleotide sequences overlap with any known deletion regions. In contrast, the FAM-labeled EGFR-19del probe was designed to fully or partially overlap with all known deletion regions. Consequently, the EGFR-WT probe serves as a positive control for the PCR assay, while the EGFR-19del probe acts as an indicator for the presence of deletion mutations. The EGFR mutation detection protocol involved preparing a 30 μL reaction mixture containing 7.5 μL PCR SuperMix (Reagent A), 7.5 μL primer-probe mixture (Reagent B), and 15 μL DNA template.

**Table 1.**
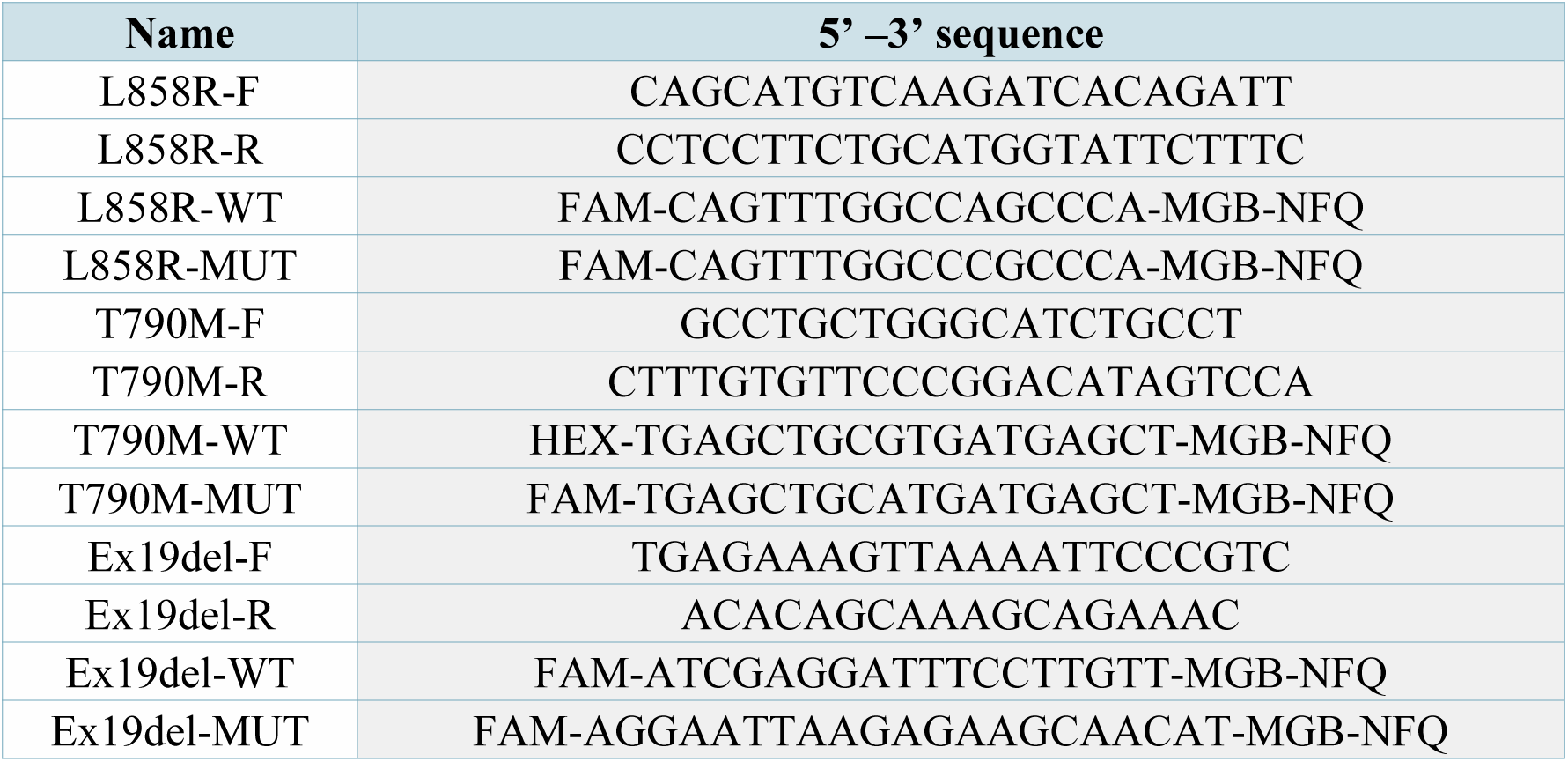
Oligonucleotide primers and probes for droplet digital polymerase chain reaction.

The ddPCR analyses were conducted using the D50 Automatic Digital PCR system (TargetingOne, Beijing, China) as follows: 30 µL of reaction mixture and 40 µL of sealing oil were added to the microfluidic chip. Subsequently, droplet generation, PCR, and droplet detection were carried out following the instructions provided with the automatic digital PCR system. Each sample generated approximately 50,000-60,000 droplets. The PCR program started with a predenaturation step at 95℃ for 10 minutes, followed by 40 amplification cycles of denaturation at 95°C for 30 seconds and annealing at 60°C for 1 minute.

## Results

### 1. Baseline Characteristics

The study cohort comprised 789 non-small cell lung cancer (NSCLC) patients with NGS confirmed EGFR mutations, including 305 cases with L858R substitution, 310 with Ex19del, and 175 with T790M mutation .One case demonstrated co-occurrence of T790M and L858R, and the exact co-mutation status beyond this individual case was not determined. The demographic and clinicopathological characteristics are summarized in **Table 3**. The cohort had a median age of 63 years (range: 30-92) and was predominantly female (61.7%, 487/789). Age distribution analysis revealed that most patients were over 60 years old (64.0%, 505/789), while only 1.5% (12/789) were younger than 40 years. The majority of patients (81.5%, 643/789) provided a single sample for analysis, with 145 patients (18.4%) contributing two samples and one patient (0.1%) providing three samples.

**Table 2.** Clinicopathological Characteristics of the Enrolled Patient Cohort.

| Parameter | Category | Number | Percentage (%) |
| --- | --- | --- | --- |
| Sex | Male | 302 | 38.3 |
|  | Female | 487 | 61.7 |
| Age (years) | <40 | 12 | 1.5 |
|  | 40-60 | 272 | 34.5 |
|  | >60 | 505 | 64 |
| Number of Samples | 1 | 643 | 81.5 |
|  | 2 | 145 | 18.4 |
|  | 3 | 1 | 0.1 |
| EGFR Mutation | L858R | 305 | 38.7 |
|  | Ex19del | 310 | 39.3 |
|  | T790M | 175 | 22.2 |

**Table 3.** Analytical Performance of EGFR Mutation Detection Across Reference Standards.

| Repl<br>icate | L858<br>R-0% | L858R-0.01% | L858R<br>-0.1% | L858<br>R-1% | T790<br>M-0<br>% | T790M<br>-0.01% | T790M<br>-0.1% | T790<br>M-1<br>% | 19DE<br>L-0% | 19DEL<br>-0.01% | 19DEL<br>-0.1% | 19DE<br>L-1% |
| --- | --- | --- | --- | --- | --- | --- | --- | --- | --- | --- | --- | --- |
| 1 | 0.00% | 0.01% | 0.14% | 0.62% | 0.00% | 0.01% | 0.13% | 0.83% | 0.00% | 0.01% | 0.25% | 1.22% |
| 2 | 0.00% | 0.01% | 0.19% | 1.15% | 0.00% | 0.01% | 0.37% | 1.43% | 0.00% | 0.01% | 0.17% | 1.83% |
| 3 | 0.00% | 0.01% | 0.12% | 1.05% | 0.00% | 0.01% | 0.11% | 1.13% | 0.00% | 0.01% | 0.17% | 1.69% |
| 4 | 0.00% | 0.01% | 0.19% | 0.88% | 0.00% | 0.01% | 0.23% | 1.17% | 0.00% | 0.02% | 0.10% | 1.29% |
| 5 | 0.00% | 0.01% | 0.13% | 1.20% | 0.00% | 0.01% | 0.13% | 0.93% | 0.00% | 0.03% | 0.11% | 1.29% |
| 6 | 0.00% | 0.01% | 0.10% | 1.04% | 0.00% | 0.02% | 0.32% | 1.24% | 0.00% | 0.03% | 0.16% | 1.03% |
| 7 | 0.00% | 0.02% | 0.13% | 1.27% | 0.00% | 0.02% | 0.17% | 1.04% | 0.00% | 0.04% | 0.23% | 1.20% |
| 8 | 0.00% | 0.02% | 0.23% | 0.79% | 0.00% | 0.02% | 0.52% | 1.05% | 0.00% | 0.04% | 0.22% | 1.16% |
| 9 | 0.00% | 0.03% | 0.11% | 0.96% | 0.00% | 0.02% | 0.07% | 0.99% | 0.00% | 0.05% | 0.24% | 1.14% |
| 10 | 0.00% | 0.04% | 0.11% | 1.25% | 0.00% | 0.02% | 0.16% | 0.94% | 0.00% | 0.06% | 0.20% | 0.86% |
| Mean | 0.00% | 0.02% | 0.15% | 1.02% | 0.00% | 0.01% | 0.22% | 1.07% | 0.00% | 0.03% | 0.18% | 1.27% |
| SD | 0.00% | 0.01% | 0.04% | 0.21% | 0.00% | 0.00% | 0.14% | 0.18% | 0.00% | 0.02% | 0.05% | 0.29% |
| CV | 0.00% | 71.50% | 30.52% | 20.74% | 0.00% | 23.60% | 63.66% | 16.37% | 0.00% | 58.77% | 28.02% | 22.81% |

The comprehensive sample collection included diverse biological specimens representative of routine clinical practice. Circulating tumor DNA (ctDNA) from plasma was available in 58 cases (25 L858R, 33 Ex19del). Pre-capture NGS libraries of cfDNA samples (cfDNA-prePCR) constituted the largest subgroup (n=319; 106 L858R, 126 Ex19del, 100 T790M). Formalin-fixed paraffin-embedded (FFPE) tissue samples were obtained from 319 patients (145 L858R, 128 Ex19del, 46 T790M), with an additional 71 pre-capture NGS libraries of FFPE samples (FFPE-prePCR: 35 L858R, 35 Ex19del, 1 T790M). Fresh tumor tissue specimens were available from 102 patients (39 L858R, 44 Ex19del, 19 T790M), while 19 patients provided pre-capture NGS libraries of fresh tumor tissue DNA (ttDNA-prePCR; 7 L858R, 12 Ex19del). Pleural effusion supernatants were collected from 48 patients (11 L858R, 11 Ex19del, 26 T790M), with corresponding pre-capture NGS libraries of pleural effusion DNA (peDNA-prePCR) available from 6 patients (4 L858R, 2 Ex19del). This comprehensive sample collection represents a diverse array of biological specimens routinely encountered in clinical practice, providing a robust foundation for methodological comparisons across different sample processing workflows and mutation subtypes. The flowchart of the study is shown as **Figure 1**.

**Figure 1.**
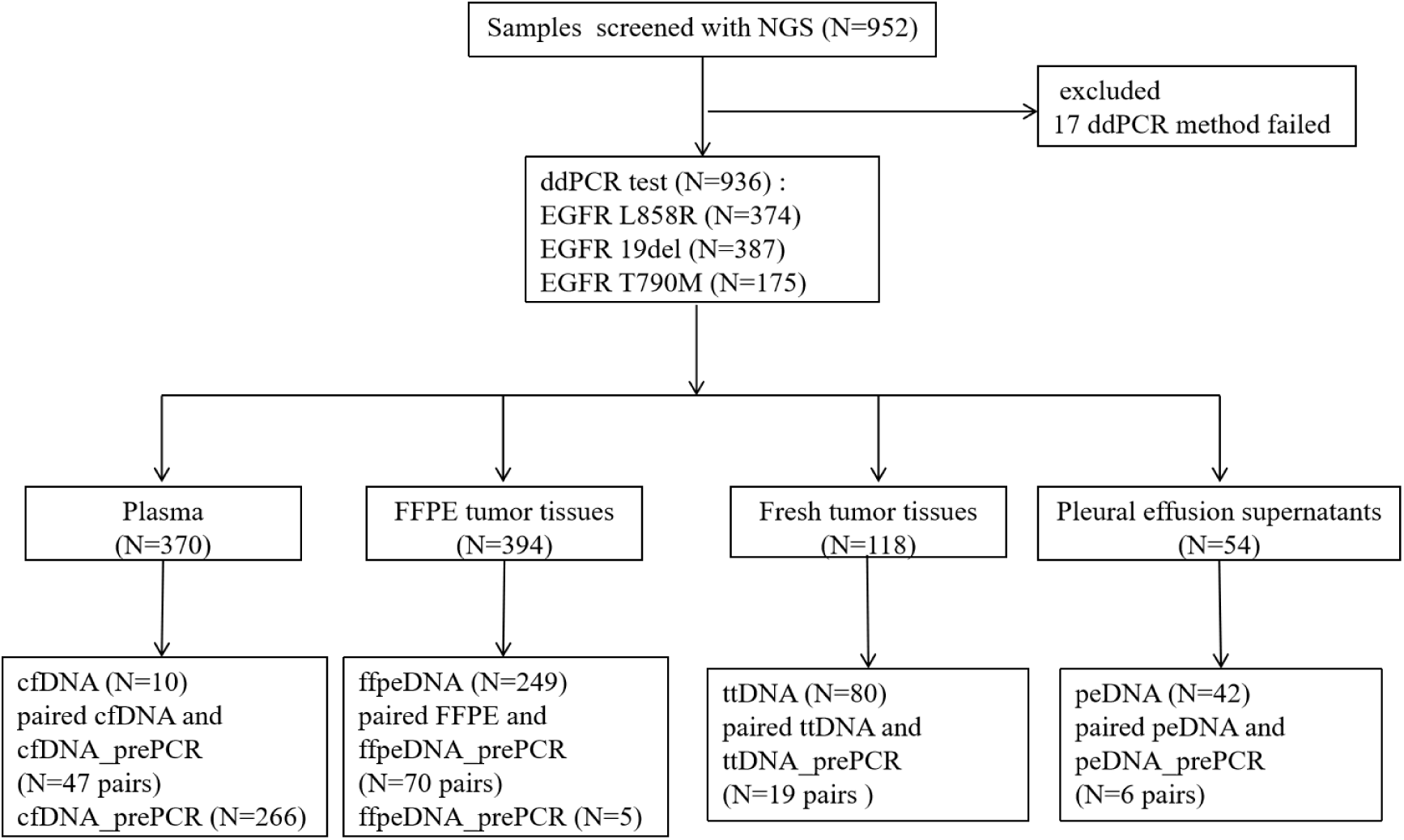
The flowchart of the study. NGS, next generation sequencing; ddPCR, digital polymerase chain reaction; cfDNA, cell free DNA; prePCR, Pre-capture NGS Library; FFPE, Formalin-Fixed Paraffin-Embedded; ttDNA, fresh tumor tissue DNA; peDNA. Pleural effusion supernatants DNA. Paired samples were compared against their source specimens to assess the consistency of the pre-capture libraries.

### 2. Determination of Limit of Detection (LOD)

The limit of detection (LOD) was established through systematic analysis of reference standards with predefined variant allele frequencies (VAFs). Given that the mass of the haploid human genome is 3.3 pg, achieving a detection limit of 0.1% requires approximately 10 ng of input DNA, consistent with the 20 ng input amount commonly employed in published studies. Accordingly, to achieve a more stringent LOD of 0.01%, approximately 100 ng of input DNA is required. Based on these calculations, we utilized 20 ng input for the 0%, 0.1%, and 1% VAF reference standards, and 100 ng input for the 0.01% VAF standard.

Each of the four reference standards for EGFR L858R, T790M, and Ex19del (0%, 0.01%, 0.1%, and 1% VAF) underwent ten replicate measurements (**Table 3**). All reference standards generated the expected results across all replicates. The measured mean VAFs for the L858R reference standards were 0.00%, 0.015%, 0.15%, and 1.02% for the 0%, 0.01%, 0.1%, and 1% standards, respectively. Similarly, the T790M standards yielded mean VAFs of 0.00%, 0.014%, 0.22%, and 1.07%, while the Ex19del standards demonstrated mean VAFs of 0.00%, 0.030%, 0.18%, and 1.27%.

For statistical validation of detection limits, 95% consistency across multiple replicates is required. All ten replicates of the 0.01% EGFR mutation reference standard with 100 ng input DNA were correctly identified as positive, confirming that sufficient input DNA is critical for avoiding false-negative results when targeting a 0.01% detection limit. These results demonstrate the robust performance of our assay across the dynamic range of clinically relevant VAFs and validate the established LOD of 0.01% for EGFR mutation detection.

### 3. ddPCR Demonstrates Robust Concordance in Mutation Detection Across All Mutation Types and Sample Sources

In the comprehensive analysis of 936 samples evaluated by both NGS and ddPCR methodologies, ddPCR exhibited excellent overall concordance for EGFR mutation detection, achieving an agreement rate of 98.72% (924/936). When using NGS as the reference standard, the positive percent agreement (PPA) of ddPCR reached 98.93%, 99.23%, and 97.14% for L858R, Ex19del, and T790M mutations, respectively. A strong correlation was observed between variant allele frequencies (VAFs) quantified by both platforms (r = 0.975; **Figure 2A**).

**Figure 2.**
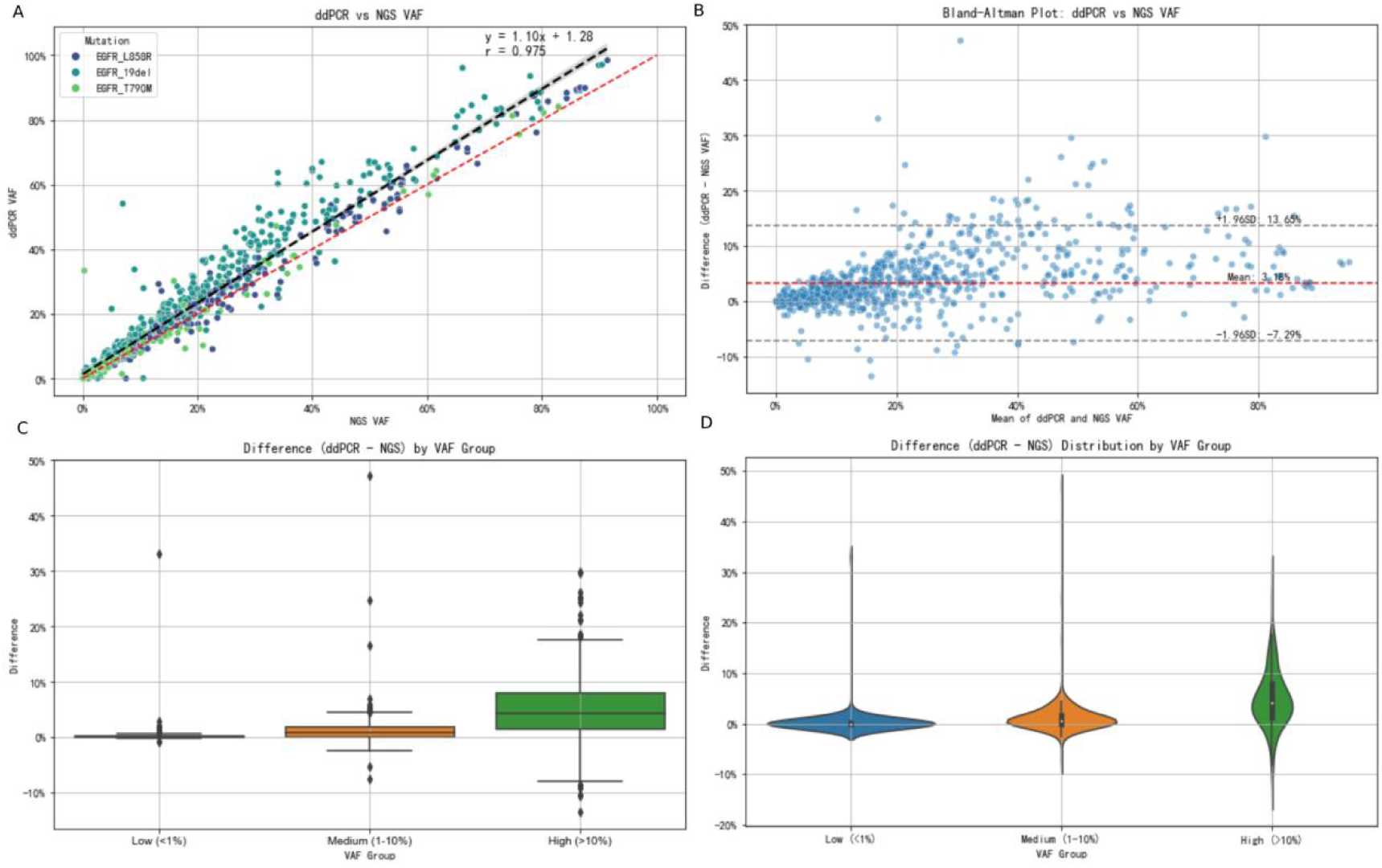
Comparison of the NGS and ddPCR tumor DNA EGFR mutation analyses. (A) Linear regression from the comparison of NGS and ddPCR VAFs (r = 0.975). Results related to the regression are presented below the graph. (B) Bland-Altman plot of the mean of NGS and ddPCR VAFs. (C) The box plots for the ddPCR and NGS VAF differences. (D)The violin plots for the ddPCR and NGS VAF differences.

Bland-Altman analysis revealed a mean bias of 3.18% (95% CI: 2.84% to 3.52%), indicating that ddPCR consistently detected approximately 3.18% higher VAF values compared to NGS. The limits of agreement (LOA) ranged from -7.29% to 13.65% (**Figure 2B**). While the 95% prediction interval suggests potential variation between individual NGS and ddPCR measurements, the overall results demonstrate significant correlation between the two methodologies.

Stratification by VAF categories (low: <1%, intermediate: 1-10%, high: ≥10%) revealed distinct patterns in methodological differences through box and violin plot visualizations (**Figure 2C, 2D**). Low VAF samples exhibited tightly clustered differences (mean: 0.03%, SD: 0.28%) with narrow graphical distributions, indicating excellent concordance for rare variant detection. Intermediate VAF samples showed moderate dispersion (mean: 0.75%, SD: 1.66%), while high VAF samples demonstrated substantial scatter (mean: 5.19%, SD: 6.26%) with extended graphical ranges. These findings indicate that both platforms demonstrate high reliability for low-frequency variant detection, critical for minimal residual disease monitoring, and methodological differences escalate with increasing VAF, with ddPCR consistently reporting higher values, potentially reflecting fundamental differences in quantification principles between digital PCR and sequencing-based approaches. Samples exhibiting high variant allele frequencies (VAFs ≥ 10%) predominantly originated from primary tumor sites. The observed mean VAF discrepancy of 5.19% between ddPCR and NGS, while statistically significant, remains within clinically acceptable limits. This margin is inconsequential for clinical decision-making, as exemplified by the 1% VAF threshold commonly used for TKI dose selection.

In summary, ddPCR demonstrated high concordance (PPA >97%) in validating NGS-detected EGFR mutations across all variant types and abundance levels. These findings support the utility of ddPCR as a reliable technique for confirmatory testing and precise quantification of EGFR mutations in clinical specimens.

### 4. Stratified Concordance Analysis by Mutation Type

For EGFR L858R point mutations (n=374), the two platforms demonstrated near-perfect correlation (r = 0.992, P < 0.001, Figure 3A) with a coefficient of determination (R²) of 0.977, indicating that 97.7% of the variance in NGS measurements was explained by ddPCR values. The mean absolute error (MAE) was 2.24%, reflecting excellent measurement precision. Bland-Altman analysis provided further insights into methodological agreement patterns across mutation subtypes. For EGFR L858R mutations, the mean bias was 1.50% with 95% limits of agreement (LoA) ranging from -4.24% to 7.23%. The 95% confidence interval (CI) for the mean difference was 1.20% to 1.80% (Figure 3B).

**Figure 3.**
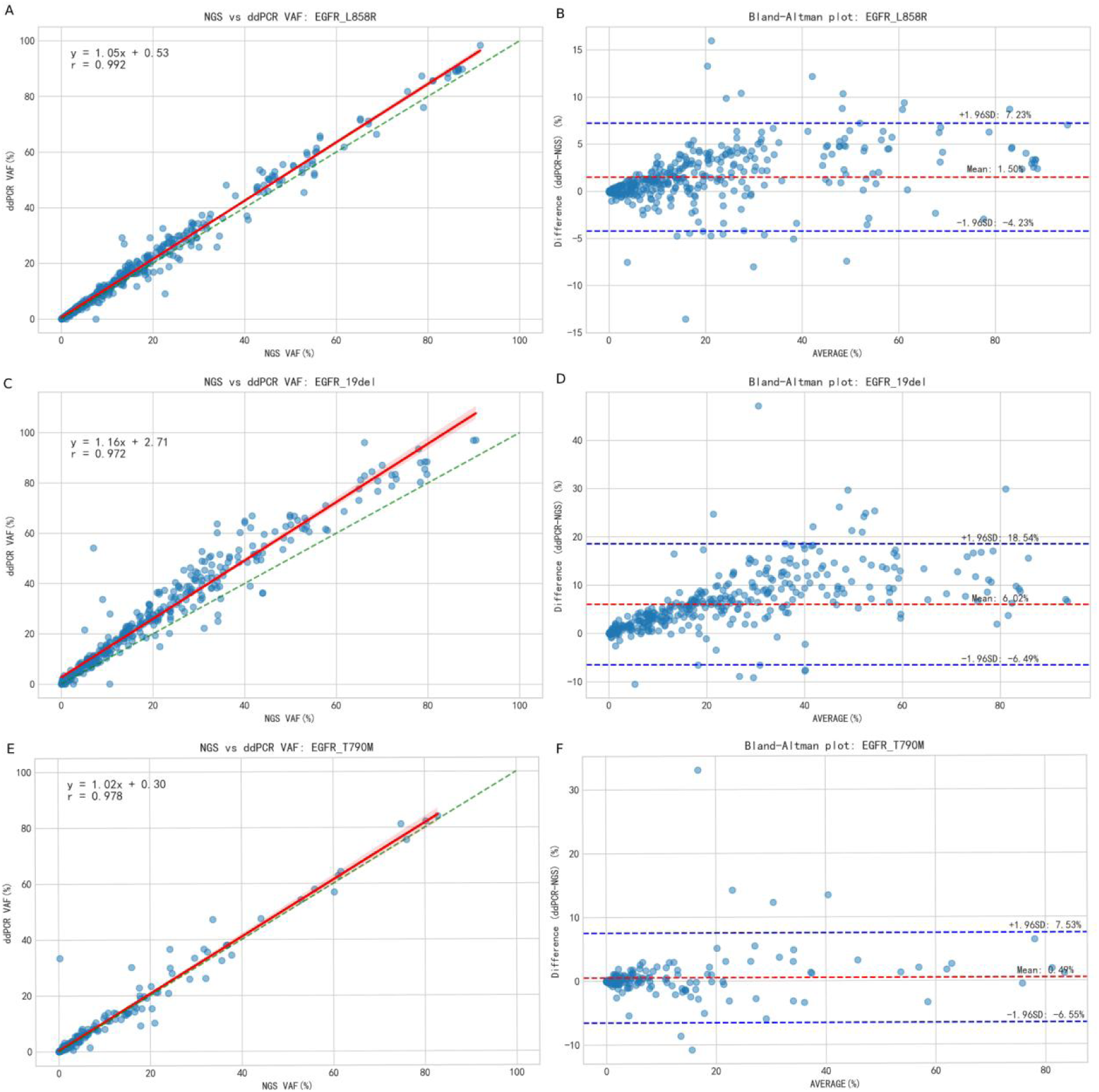
Comparison of the NGS and ddPCR tumor DNA EGFR mutation analyses. (A) Linear regression from the comparison of NGS and ddPCR VAFs (r = 0.992) with EGFR L858R mutation. (B) Bland-Altman plot of the mean of NGS and ddPCR VAFs with EGFR L858R mutation. (C) Linear regression from the comparison of NGS and ddPCR VAFs (r = 0.972) with EGFR Ex19del mutation. (D) Bland-Altman plot of the mean of NGS and ddPCR VAFs with EGFR Ex19del mutation. (E) Linear regression from the comparison of NGS and ddPCR VAFs (r = 0.978) with EGFR T790M mutation. (F) Bland-Altman plot of the mean of NGS and ddPCR VAFs with EGFR T790M mutation.

Ex19del (n=387) showed strong correlation (r = 0.972, P < 0.001, Figure 3C) with an R² of 0.860, though with moderately higher absolute error (MAE = 6.40%) compared to other mutation types. Bland-Altman analysis showed EGFR Ex19del variants demonstrated a substantially higher mean bias of 6.02% with wider limits of agreement (-6.51% to 18.55%) (Figure 3D) and a CI of 5.38% to 6.66%.

Notably, the EGFR T790M resistance mutation cohort (n=175) exhibited exceptional methodological agreement (r = 0.978, P < 0.001; R² = 0.955, Figure 3E) and achieved the lowest mean absolute error (MAE = 1.55%) among all mutation subtypes analyzed. Bland-Altman analysis of the EGFR T790M cohort showed the most favorable agreement profile, with a minimal mean bias of 0.49% and narrow limits of agreement (-6.57% to 7.55%) (Figure 3F). Importantly, the 95% CI for the mean difference in T790M measurements included zero (-0.05% to 1.03%), indicating no statistically significant systematic bias between the two methodologies for this mutation type.

This comprehensive stratified analysis revealed high methodological concordance between ddPCR and NGS across all major EGFR mutation subtypes, with performance characteristics varying by specific mutation type. These findings demonstrate mutation-dependent performance characteristics, with T790M detection showing superior agreement between platforms, while Ex19del exhibit greater methodological variability. The consistent high correlation across all subtypes nevertheless supports the reliability of ddPCR for EGFR mutation quantification in clinical practice.

### 5. Consistency analysis grouped by sample type

Method comparison analysis revealed strong concordance between ddPCR and NGS across all specimen types. For plasma-derived circulating tumor DNA (cfDNA, n=57), excellent correlation was observed (r=0.958, P<0.001; R²=0.877, Figure 4A) with a mean absolute error (MAE) of 5.66%. Bland-Altman analysis revealed systematic differences in methodological agreement across various specimen types. For plasma circulating tumor DNA (cfDNA, n=57), a mean bias of 4.43% was observed with wide limits of agreement (LoA) ranging from -10.97% to 19.83% (95% CI for mean bias: 2.34-6.51%) (Figure 4B). Formalin-fixed paraffin-embedded DNA (ffpeDNA, n=319) showed high methodological agreement (r=0.973, P<0.001; R²=0.915, Figure 4C) with moderate error (MAE=4.34%). Formalin-fixed paraffin-embedded DNA (ffpeDNA, n=319) showed moderate mean bias (3.44%) with LoA of -6.61% to 13.49% (Figure 4D). Tissue-derived DNA (ttdna, n=99) exhibited near-perfect correlation (r=0.981, P<0.001; R²=0.910, Figure 4E) despite slightly higher MAE (4.87%). Tissue-derived DNA (ttdna, n=99) exhibited a mean bias of 4.65% and LoA from -4.99% to 14.29% (Figure 4F). Plasma-derived extracellular DNA (pedna, n=48) demonstrated exceptional concordance (r=0.988, P<0.001; R²=0.946, Figure 4G) with the second-lowest MAE (2.35%). Notably, plasma extracellular DNA (pedna, n=48) displayed the most favorable agreement profile with minimal mean bias (2.19%) and the narrowest LoA (-4.42% to 8.81%) (Figure 4H).

**Figure 4.**
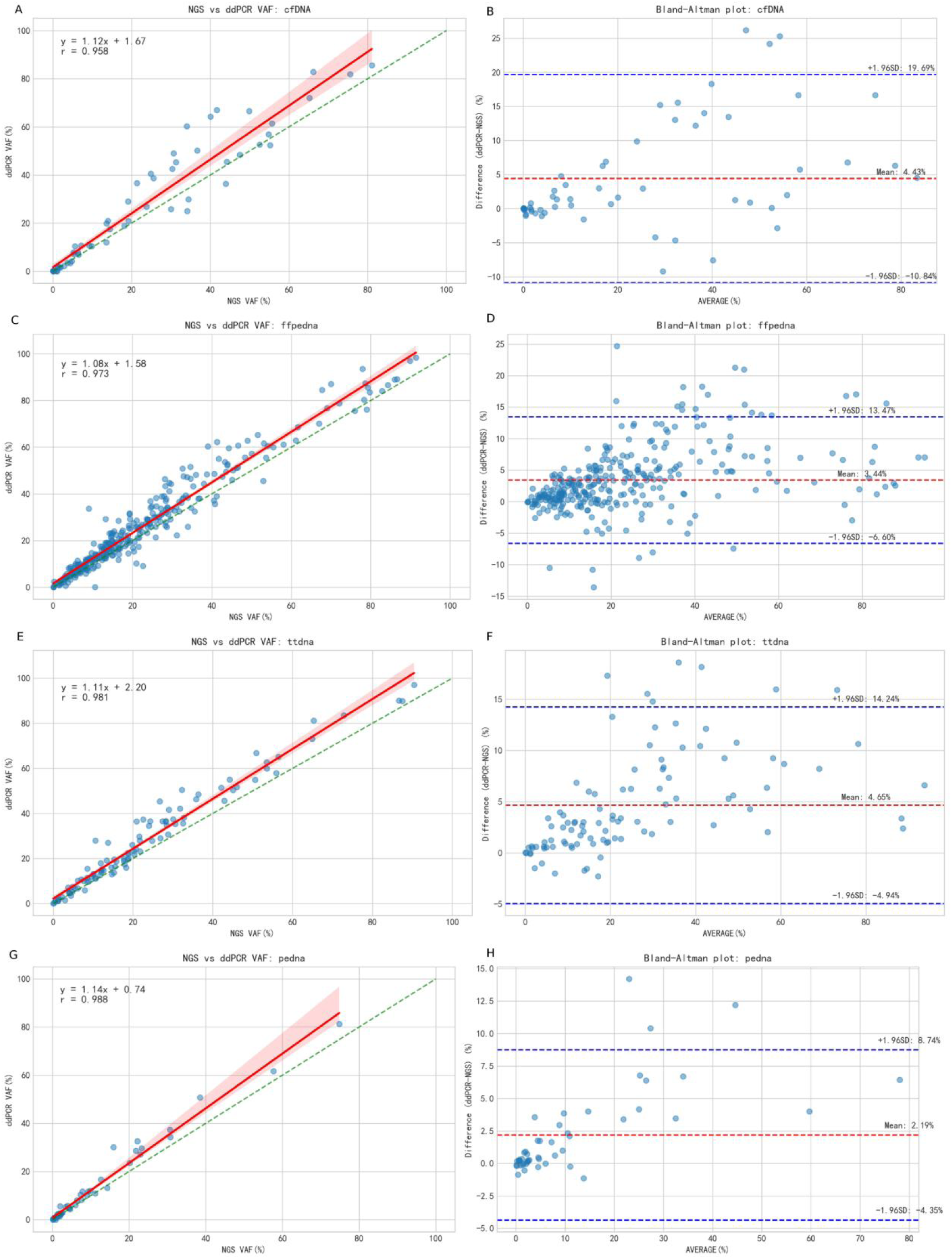
Consistency analysis of the NGS and ddPCR tumor DNA EGFR mutation grouped by sample type. (A) Linear regression from the comparison of NGS and ddPCR VAFs with cfDNA. (B) Bland-Altman plot of the mean of NGS and ddPCR VAFs with cfDNA. (C) Linear regression from the comparison of NGS and ddPCR VAFs with FFPE DNA. (D) Bland-Altman plot of the mean of NGS and ddPCR VAFs with FFPE DNA. (E) Linear regression from the comparison of NGS and ddPCR VAFs with ttDNA. (F) Bland-Altman plot of the mean of NGS and ddPCR VAFs with ttDNA. (G) Linear regression from the comparison of NGS and ddPCR VAFs with peDNA. (H) Bland-Altman plot of the mean of NGS and ddPCR VAFs with peDNA.

cfDNA-prePCR (n=313) demonstrated even stronger correlation (r=0.962, P<0.001; R²=0.908, Figure 5A) and improved accuracy (MAE=2.54%). cfDNA-prePCR (n=313) demonstrated significantly improved agreement, with reduced mean bias (1.91%) and narrower LoA (-8.73% to 12.55%) (Figure 5B). ffpeDNA-prePCR (n=75) and ttdna-prePCR (n=19) both maintained excellent correlation coefficients (r=0.990 and r=0.992, respectively; both P<0.001) (Figure 5C, 5E) with moderate measurement errors (MAE=4.68% and 5.76%, respectively). ffpeDNA-prePCR (n=75) and ttdna-prePCR (n=19) demonstrated mean biases of 4.36% and 5.71% (Figure 5D, 5F), respectively. Pedna-prePCR (n=6) showed perfect correlation (r=0.999, P<0.001, Figure 5G) though with limited sample size. Among pre-PCR processed samples, pedna-prePCR (n=6) showed substantial mean bias of 4.56% (Figure 5H) with wide confidence intervals due to limited sample size.

**Figure 5.**
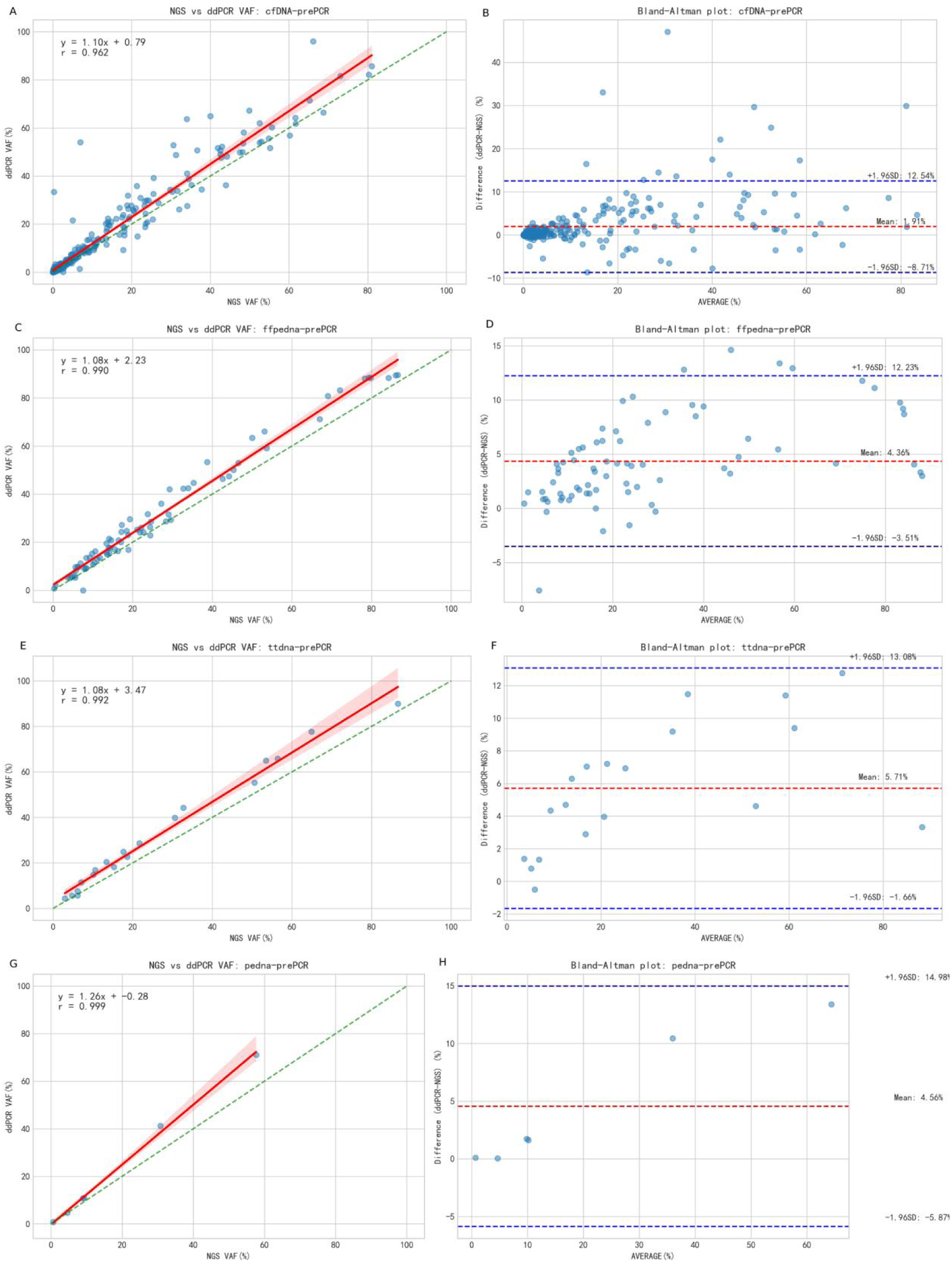
Consistency analysis of the NGS and ddPCR tumor DNA EGFR mutation grouped by sample type. (A) Linear regression from the comparison of NGS and ddPCR VAFs with cfDNA-prePCR. (B) Bland-Altman plot of the mean of NGS and ddPCR VAFs with cfDNA-prePCR. (C) Linear regression from the comparison of NGS and ddPCR VAFs with ffpeDNA-prePCR. (D) Bland-Altman plot of the mean of NGS and ddPCR VAFs with ffpeDNA-prePCR. (E) Linear regression from the comparison of NGS and ddPCR VAFs with ttDNA-prePCR. (F) Bland-Altman plot of the mean of NGS and ddPCR VAFs with ttDNA-prePCR. (G) Linear regression from the comparison of NGS and ddPCR VAFs with peDNA-prePCR. (H) Bland-Altman plot of the mean of NGS and ddPCR VAFs with peDNA-prePCR.

### 6. Quantitative Evidence Supporting Pre-capture NGS Library Fidelity

Analysis of 44 paired ctDNA versus Library Preparations samples demonstrated outstanding concordance between native plasma-derived circulating tumor DNA (ctDNA) and library-processed specimens across both major EGFR mutation types. The methodological comparison showed a strong correlation (r = 0.993, P < 0.001, Figure 6A) with an R² of 0.986, while maintaining a low MAE of 1.65%. These results collectively indicate that library preparation introduces minimal additional variability in ddPCR measurements. Bland-Altman analysis was employed to quantitatively assess the agreement between different sample processing methodologies across various EGFR mutation types. The comparison between native ctDNA and library-processed samples across both mutation types (n=44) demonstrated intermediate performance, with a mean bias of 1.65% (95% CI: 1.21-2.09%) and limits of agreement from -3.13% to 6.43% (Figure 6B).

**Figure 6.**
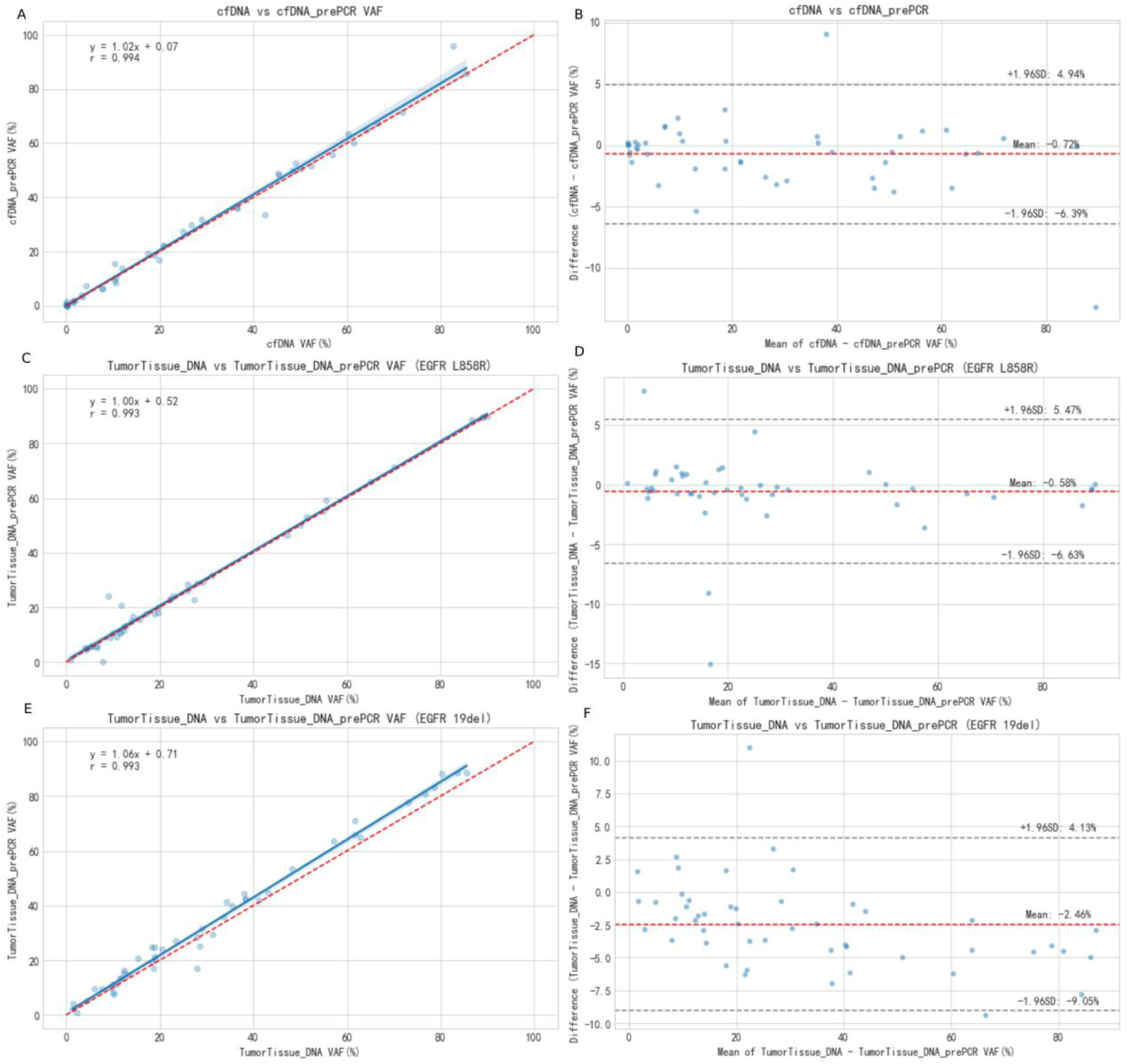
Comparison of the tumor DNA and paired Pre-capture NGS Library analyses. (A) Linear regression from the comparison of cfDNA and cfDNA-prePCR VAFs. (B) Bland-Altman plot of the mean of cfDNA and cfDNA-prePCR VAFs. (C) Linear regression from the comparison of tumor tissue DNA and tumor tissue DNA prePCR VAFs with EGFR L858R mutation. (D) Bland-Altman plot of the mean of tumor tissue DNA and tumor tissue DNA prePCR VAFs with EGFR L858R mutation. (E) Linear regression from the comparison of tumor tissue DNA and tumor tissue DNA prePCR VAFs with EGFR Ex19del mutation. (F) Bland-Altman plot of the mean of tumor tissue DNA and tumor tissue DNA prePCR VAFs with EGFR Ex19del mutation.

Evaluation of 47 paired tumor tissue DNA versus Library Preparations with EGFR L858R mutation samples showed exceptional methodological concordance for EGFR L858R point mutations. The analytical comparison demonstrated an almost perfect correlation (r = 0.998, P < 0.001, Figure 6C) with an R² value of 0.996, revealing that 99.6% of the variability in library measurements was captured by native FFPE DNA values. The exceptional measurement precision was further corroborated by an exceptionally low MAE of 1.08%. In the case of EGFR L858R mutations (n=47), superior agreement was observed with a mean bias of 1.08% (95% CI: 0.74-1.42%) and narrower limits of agreement (-2.69% to 4.85%) (Figure 6D).

Analysis of 49 paired tumor tissue DNA versus Library Preparations with EGFR Ex19del specimens demonstrated excellent concordance between ddPCR measurements from native FFPE DNA and corresponding library preparations for EGFR Ex19del variants. The methodological comparison revealed a near-perfect Pearson correlation coefficient of 0.991 (P < 0.001,, Figure 6E), with a coefficient of determination (R²) of 0.982, indicating that 98.2% of the variance in library measurements could be explained by native FFPE DNA values. The mean absolute error (MAE) was 2.07%, reflecting minimal practical difference between the two sample processing methodologies in clinical measurement. For EGFR Ex19del mutations in FFPE DNA versus library preparations (n=49), the mean bias was 2.07% (95% CI: 1.58-2.56%) with limits of agreement ranging from -3.89% to 8.03% (Figure 6F).

All correlation analyses demonstrated statistical significance (P < 0.001), confirming that the observed methodological relationships were not attributable to random chance. Rigorous quality control measures were systematically implemented throughout the study, including exclusion of samples with incomplete paired measurements, to ensure analytical reliability and result validity.

## Discussion

In this study, we established three landmark contributions to molecular diagnostics in NSCLC through its large-scale, multi-sample and multi-mutation design (n=936). The comprehensive validation across plasma, FFPE, fresh tissue, and pleural effusion specimens provides unprecedented evidence for implementing NGS as a gold-standard detection platform across diverse clinical scenarios. This large-scale comparative analysis provides robust evidence supporting the high concordance between NGS and ddPCR for detecting key EGFR mutations (L858R, Ex19del, T790M) and quantifying their VAF across diverse sample types relevant to NSCLC clinical management.

We found ddPCR demonstrated high concordance (PPA, 98.72%) in validating NGS-detected EGFR mutations across all variant types and abundance levels. This high level of agreement across nearly 1000 mutation-positive samples, encompassing critical variants like L858R (98.93%), Ex19del (99.23%), and T790M (97.14%), strongly reinforces the reliability of both platforms. The overall Pearson correlation coefficient was 0.975 (P < 0.001), indicating a very strong positive relationship between the two platforms. The mean absolute error (MAE) was 3.83%, with a mean difference of 3.18% (95% CI: 2.84% to 3.52%) and standard deviation of differences of 5.34%. These metrics collectively demonstrate excellent methodological agreement across the large sample cohort. The overall mutation detection concordance rate of 0.987 significantly surpasses rates (0.92–0.95) previously reported in smaller single-center studies. This enhanced performance likely reflects both the statistical power of our cohort size and the inclusion of diverse sample types.

These findings support the utility of ddPCR as a reliable technique for confirmatory testing and precise quantification of EGFR mutations in clinical specimens. It validates their established roles in clinical practice and underscores that either method can provide clinically actionable results for these common mutations. This concordance is particularly reassuring given the distinct technological principles underlying NGS (amplification and sequencing) and ddPCR (endpoint partitioning and fluorescence detection), mitigating concerns about systematic biases inherent to either platform alone.

Despite the high overall concordance, 12 discordant samples (1.28%) were identified, comprising 4 L858R, 3 Ex19del, and 5 T790M mutations. Comprehensive analysis revealed several potential contributing factors. Pre-PCR processed samples accounted for 7 of 12 discordant cases (58.3%), including 6 cfDNA-prePCR and 1 FFPE-prePCR specimens. Therefore, the primary risk factor may lie in the pre-amplification Artifacts. The remaining 5 discordant cases included pleural effusion DNA (2 cases), conventional cfDNA (2 cases), and FFPE DNA (1 case). These sample types present inherent challenges, such as low DNA quantity and quality, fragmentation and degradation and PCR inhibitors. The observed discordances primarily originate from sample pre-processing methodologies (particularly pre-amplification) and intrinsic sample quality limitations rather than fundamental incompatibility between ddPCR and NGS technologies. In addition, the negative results of digital PCR for these samples may also be attributed to degradation caused by repeated freezing and thawing or the mutation types not being covered by the digital PCR detection range.

Beyond mere detection, the high correlation in VAF quantification is crucial for clinical interpretation. VAF is increasingly recognized as a potential biomarker for tumor burden, clonal dynamics, and response assessment. The strong VAF correlation observed suggests that both methods provide comparable estimates of mutation abundance, facilitating consistent longitudinal monitoring regardless of the platform chosen for initial testing or follow-up.

The comparative analysis demonstrated high concordance between ddPCR and NGS across all EGFR mutation subtypes. For EGFR_L858R mutations (n=374), we observed near-perfect correlation (r=0.9919, P<0.001; R²=0.9769) with minimal measurement discrepancy (MAE=2.24%). Similarly, EGFR_ Ex19del variants (n=387) showed strong correlation (r=0.9717, P<0.001; R²=0.8604) with moderate absolute error (MAE=6.40%). The EGFR_T790M cohort (n=175) exhibited excellent methodological agreement (r=0.9777, P<0.001; R²=0.9547) and the lowest mean absolute error (MAE=1.55%) among all mutation types analyzed. The Bland-Altman analysis revealed differential agreement patterns across EGFR mutation subtypes . EGFR_L858R mutations (n=374) showed a mean bias of 1.50% (95% CI: 1.20-1.80%) with limits of agreement from -4.24% to 7.23%. EGFR_ Ex19del variants (n=387) exhibited substantially greater mean bias (6.02%; 95% CI: 5.38-6.66%) and wider limits of agreement (-6.51% to 18.55%). In contrast, EGFR_T790M mutations (n=175) demonstrated the highest level of agreement, with a mean bias of 0.49% (95% CI: -0.05-1.03%) and limits of agreement ranging from -6.57% to 7.55%. These findings demonstrate mutation-dependent performance characteristics, with T790M detection showing superior agreement between platforms, while Ex19del exhibit greater methodological variability. The consistent high correlation across all subtypes nevertheless supports the reliability of ddPCR for EGFR mutation quantification in clinical practice.

The analytical performance between ddPCR and NGS was evaluated across multiple specimen types. All sample types demonstrated strong correlation (r=0.958-0.999, all P<0.001) and high coefficients of determination (R²=0.877-0.946). The lowest measurement errors were observed in cfDNA (MAE=2.54%) and pedna samples (MAE=2.35%), while the highest errors were noted in ttdna-prePCR (MAE=5.76%) and standard cfDNA samples (MAE=5.66%). Pre-PCR processing generally improved correlation metrics in cfDNA and pedna specimens compared to their standard counterparts. The analysis demonstrates that pre-analytical processing influences methodological agreement, with pre-PCR samples generally showing improved correlation coefficients. Notably, plasma-derived DNA (both cfDNA and pedna) exhibited the lowest measurement errors following pre-PCR processing, suggesting potential benefits of this preparatory step for liquid biopsy applications.

A particularly noteworthy and clinically practical finding is the remarkable fidelity of pre-capture NGS libraries to their original source materials. The near-perfect VAF concordance (0.993 for cfDNA libraries vs. cfDNA; 0.998, and 0.991 for tumor tissue libraries vs. tumor DNA with EGFR L858R, and Ex19del, respectively) demonstrates that library preparation introduces minimal additional variability in ddPCR measurements, and these libraries accurately preserve the mutational information present in the primary sample. This has profound implications for resource utilization and retrospective analysis. When the original source material (e.g., precious biopsy DNA or limited cfDNA) is depleted or unavailable, pre-capture libraries represent a reliable alternative for additional testing, including validation by ddPCR, reflex testing for other targets, or future investigations using emerging technologies. This maximizes the utility of scarce patient samples and reduces the need for repeat invasive procedures.

The comprehensive analysis revealed distinct patterns of methodological agreement across different experimental conditions. EGFR L858R mutations demonstrated the highest level of methodological consistency, as evidenced by the smallest mean bias (1.08%) and the narrowest limits of agreement among all comparison groups. In contrast, EGFR Ex19del variants showed relatively reduced agreement, with both higher mean bias (2.07%) and wider limits of agreement. The comparison between native ctDNA and library samples exhibited intermediate performance, suggesting that library preparation introduces measurable but clinically acceptable variability in the detection process.

Notably, the 95% confidence intervals for the mean bias excluded zero in all three comparison groups, indicating statistically significant systematic differences between the measurement methodologies. However, these systematic biases, while statistically significant, remain within clinically acceptable ranges for most diagnostic applications, thereby supporting the robustness of pre-capture NGS library preparation methodologies.

The high inter-method concordance supports complementary deployment of NGS and ddPCR. NGS remains optimal for initial comprehensive profiling due to its multiplexing capability. NGS can explore a broad spectrum of genetic aberrations in a single run, and the possibilities to detect translocations and amplifications are expanding quickly. Thus, with the expanding knowledge of resistance mechanisms and possible targeted treatments (in development) for these, the detection of a broad set of genetic aberrations seems desirable. While ddPCR’s superior sensitivity and precision suit minimal residual disease (MRD) monitoring, early detection of resistance mutations (e.g., T790M), and VAF-guided dose optimization. Additionally, ddPCR offers two notable advantages: first, a fast turnaround time — it can typically deliver results to clinicians within one working day when urgency is required; second, a significant cost advantage — it can substantially reduce labor costs, operational costs, and expenses for reagents and consumables. These costs are often driven by complex operational and analytical processes as well as long experimental cycles, and ddPCR ultimately enables high-frequency detection of specific biomarkers at a lower cost. This study validates integrated workflows (e.g., NGS screening with ddPCR confirmation/monitoring). Pre-capture libraries maximize biospecimen utility—particularly relevant for the 20–30% of NSCLC patients with insufficient tumor tissue. For advanced-stage patients requiring repeated sampling, this approach significantly enhances diagnostic feasibility.

These results provide a solid foundation for developing integrated diagnostic paradigms. The high concordance supports the strategic use of both NGS and ddPCR. The data confirm that the test results of the two technologies are reliably comparable. The reliability of pre-capture libraries offers a practical solution for maximizing information from limited samples, enabling reflex testing without consuming irreplaceable source material. This study contributes valuable data towards standardizing cross-platform validation and result interpretation for EGFR testing. Future research should explore the concordance for less common EGFR mutations and other actionable genes, investigate the impact of very low VAF levels (<1%) on concordance, and prospectively evaluate the clinical utility of using pre-capture libraries for specific applications like therapy response monitoring.

However, it is important to acknowledge limitations. A key limitation of the current performance validation is the lack of NGS-negative and healthy control samples, which prevents the assessment of specificity and negative percent agreement (NPA). To address this, future work must include the assembly of a negative control cohort. This cohort should consist of samples confirmed to be negative by NGS, which must then be tested via ddPCR to determine whether they yield negative results, thereby enabling the calculation of specificity and NPA. Besides, the study focused on three common EGFR mutations, while concordance for rarer mutations or other genes might differ. While the sample size is large, the analysis was retrospective. Prospective validation in specific clinical scenarios (e.g., monitoring minimal residual disease) would further solidify these findings.

## Conclusion

In conclusion, this large scale study definitively demonstrates high concordance between NGS and ddPCR for detecting major EGFR mutations and quantifying VAF across diverse NSCLC sample types. Crucially, it establishes pre-capture NGS libraries as faithful representations of original samples, unlocking significant potential for efficient resource utilization and flexible testing strategies. These findings empower clinicians and laboratories to confidently leverage the complementary strengths of NGS and ddPCR, paving the way for more robust, adaptable, and patient-centric molecular testing pathways in NSCLC. The demonstrated concordance and reliability of pre-capture libraries will accelerate the transition from single-method dependency to integrated precision diagnostics, ultimately advancing personalized therapeutic decision-making.

## Notes

### Competing Interest Statement

The authors have declared no competing interest.

### Funding

- Shenzhen Science and Technology Innovation Committee Technical Research Project: JSGG20201103153801005

